# Accurately estimating pathway activity in single cells for clustering and differential analysis

**DOI:** 10.1101/2023.03.27.534310

**Authors:** Daniel Davis, Avishai Wizel, Yotam Drier

## Abstract

Inferring which and how biological pathways and gene sets are changing is a key question in many studies that utilize single-cell RNA sequencing. Typically, these questions are addressed by quantifying the enrichment of known gene sets in lists of genes derived from global analysis. Here we offer SiPSiC, a new method to infer pathway activity in each cell. This allows more sensitive differential analysis and utilizing pathway scores to cluster cells and compute UMAP or other similar projections. We apply our method on datasets of COVID-19, lung adenocarcinoma and glioma, and demonstrate its utility. SiPSiC analysis is consistent with findings reported by previous analyses in many cases, but also reveals the differential activity of novel pathways, enabling us to suggest new mechanisms underlying the pathophysiology of these diseases and demonstrating SiPSiC’s high accuracy and sensitivity in detecting biological function and traits. In addition, we demonstrate how it can be used to better classify cells based on activity of biological pathways instead of single genes and its ability to overcome patient specific artifacts.

## Introduction

Single-cell RNA sequencing (scRNA-seq) has become a staple technique in biomedical research, allowing for a deeper understanding of tissue characteristics and heterogeneity in health and disease (Chi & Deng, 2020). Pathway analysis serves as a crucial tool for interpreting gene expression data, particularly in the context of scRNA-seq. Typically, this is accomplished by identifying a set of interesting genes, such as differentially expressed genes, and examining their enrichment within biological pathways. While such approaches are very useful, there are many potential benefits to estimate pathway activity in each cell first, and only then utilize this information for downstream analysis. This allows both to overcome limitations of single-cell data, such as inaccurate estimation of the expression of a single gene in a single cell, and to utilize pathway activity for unsupervised analysis such as clustering.

Although methods for pathway analysis in bulk RNA-seq have been developed (Drier et al., 2013; Hänzelmann et al., 2013; H. Wang et al., 2016), they often prove inadequate for scRNA-seq data due to its unique characteristics (Noureen et al., 2022). Past efforts to estimate pathway activity in single cells have utilized techniques such as AUCell (Aibar et al., 2017) and single sample gene set enrichment analysis (ssGSEA) (Barbie et al., 2009). However, these methods were not designed for this type of analysis and are not always optimal.

Here we introduce Single Pathway analysis in Single Cells (SiPSiC), a new method tailored for analyzing the activity of a gene set, or biological pathway, in single cells. SiPSiC reaches high sensitivity by relying on the normalized expression of all genes, weighted by their relative rank. Using SiPSiC, we reanalyze scRNA-seq data of COVID-19, lung adenocarcinoma and glioma, identify both known and novel cellular pathways involved in these diseases, and demonstrate SiPSiC’s high accuracy and superior ability to identify changes in pathway activity missed by the original analyses. Furthermore, we propose new approaches for data clustering and visualization based on SiPSiC scores, mitigating biases inherent in scRNA-seq data and emphasizing functional similarity based on shared biological processes. Through comparative analyses with existing methods, we showcase SiPSiC’s superior accuracy, sensitivity, and efficiency, positioning it as a valuable tool for pathway analysis in single-cell studies.

## Results

### A new tool to infer pathway activity in single cells

Single-cell RNA sequencing (scRNA-seq) data often suffers from sparsity due to high dropout rates, posing challenges for accurate pathway analysis. To address this issue, we introduce SiPSiC, a new tool designed to calculate pathway scores for each individual cell and each gene set by using gene expression values normalized by the expression levels in other cells and weighted by the rank of the average gene expression across all genes of the gene set (see Methods). This robust weighted normalization enables accurate estimation of pathway activity, even in small datasets or when pathway gene coverage is limited.

Available on both GitHub and Bioconductor, SiPSiC is an easily installed and well documented tool to dissect single cell level differences, allowing its users to interrogate tissue physiology and heterogeneity with high sensitivity and accuracy.

### SiPSiC reveals differential activity of pathways with potential therapeutic implications in SARS-CoV-2 infected cells

We applied SiPSiC to investigate changes in activity of biological pathways after SARS-CoV-2 infection in two distinct datasets: single nuclei RNA-seq data from recently deceased COVID-19 patients (Melms et al., 2021) and scRNA-seq of African green monkeys infected with SARS-CoV-2 or inactivated virus (Speranza et al., 2021). The original analysis of the human data included gene-based clustering from which cluster markers were inferred to elucidate cellular response to SARS-CoV-2 infection. The monkey data was originally analyzed by principal component analysis (PCA), clustering and Fast Gene Set Enrichment Analysis (Korotkevich et al., 2021). We calculated pathway scores per cell for each of the 50 MSigDB hallmark gene sets (Liberzon et al., 2015), and compared pathway scores between SARS-CoV-2 positive and negative controls. Our findings can be largely divided into three categories: 1) Innate immune response of pneumocytes; 2) Pathways modulated by the virus to support its life cycle; and 3) Adaptive immune response in B and CD8+ T cells.

First, we applied SiPSiC to human alveolar cells (of both type 1 and 2; 4575 cells from COVID-19 patients vs 4303 control cells). 12 out of 50 pathways were downregulated in COVID-19 patients, and 31 upregulated (Student’s t-test, FDR < 0.01, Figure 1A; see Supplemental Table S1). In addition to interferon response which was also reported as upregulated in alveolar type 2 (AT2) cells by Melms et al., we identified many more upregulated pathways involved in innate immune response and its implications, among them are genes involved in the complement pathway, DNA repair and Wnt/beta catenin signaling. Indeed, the complement system was shown to be hyperactivated in severe SARS-CoV-2 infections (Afzali et al., 2021), SARS-CoV-2 induces DNA damage (Gioia et al., 2023), and WNT5A is upregulated in severe cases of COVID-19 (Choi et al., 2020). We also found upregulation of the mitotic spindle (FDR < 1.1*10^−4^) and E2F targets (FDR < 2.3*10^−15^), which is consistent with demonstrated hyperplasia of pneumocytes in humans and monkeys post SARS-CoV-2 infection (Ackermann et al., 2020; Speranza et al., 2021). In addition, Melms et al. found that the alveolar cells of the COVID-19 group showed lower expression of the ETV5 transcription factor required for maintaining AT2 cell identity compared to alveolar cells of the control lungs. Combined, these findings suggest that infected pneumocytes try to compensate for the damaged tissue by advancing the cell cycle and differentiating towards AT1 cells. Furthermore, SiPSiC detected upregulation of the hallmark apoptosis pathway (FDR < 1.1 ∗ 10^−30^), consistent with detection of many apoptotic cells in human airway epithelium cultures infected with SARS-CoV-2 (Zhu et al., 2020). This may relate to apoptosis induction by the viral ORF3a protein (Ren et al., 2020). Additionally, it may relate to apoptosis induction by TP53, and indeed SiPSiC also detected upregulation of the TP53 pathway (FDR < 7.9 ∗ 10^−133^). Prior research showed that viral infection upregulates TP53 by type 1 interferon signaling (Takaoka et al., 2003) and, on the other hand, that TP53 both activates the interferon pathway and promotes type 1 interferon release from cells undergoing viral infection (Muñoz-Fontela et al., 2008). These findings together suggest this positive feedback loop of interferon and TP53 may also play a role in SARS-CoV-2 infection.

**Figure 1.**
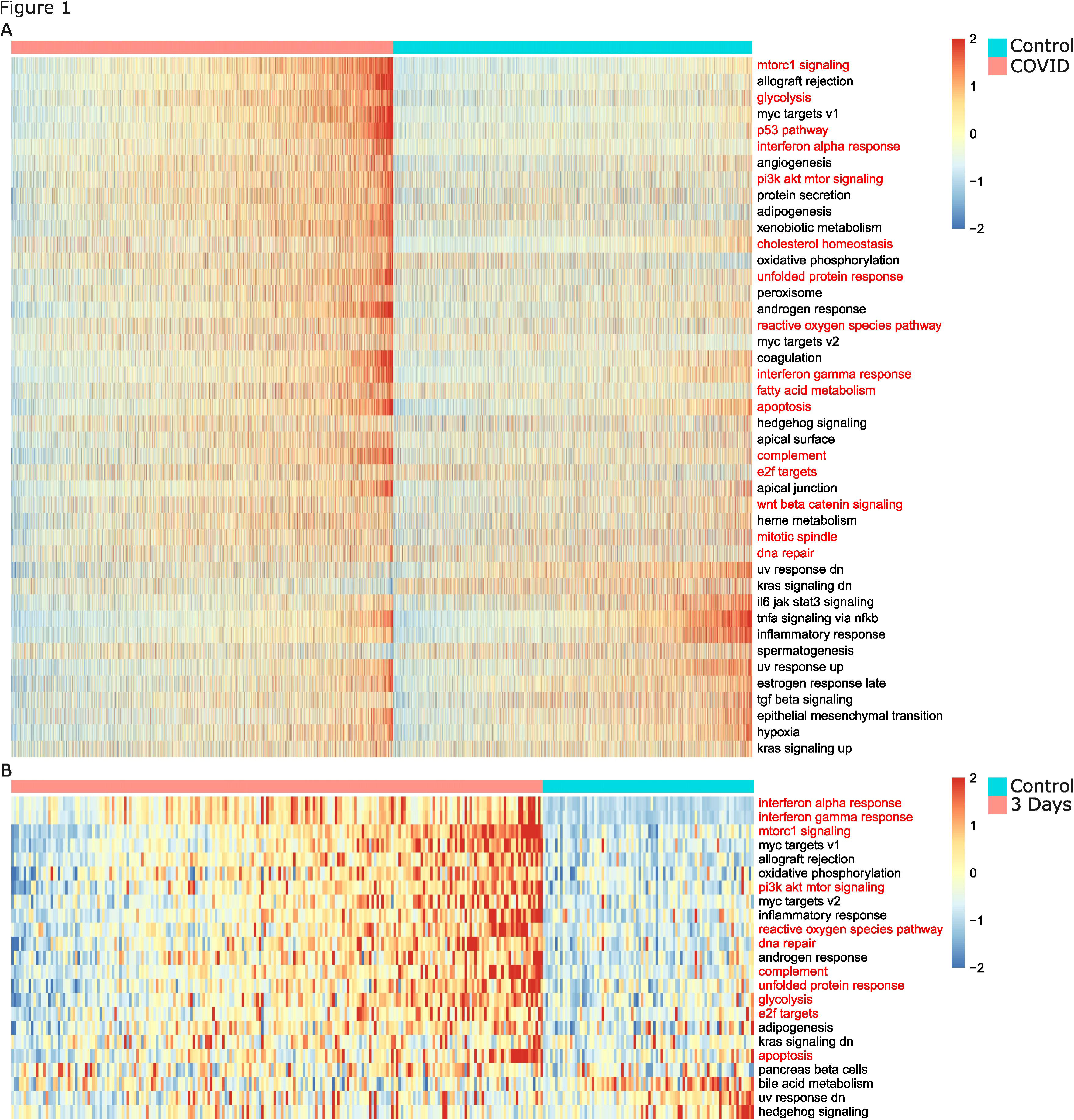
**(A-B)** Heatmaps depicting z-scores of SiPSiC scores of alveolar cells for all differential hallmark pathways (FDR < 0.01) for COVID-19 patients and the control group **(A)** or SARS-CoV-2 infected monkeys and controls **(B)**. Pathways are sorted by significance of differential scores, cells in each cell group are sorted by their average z-score across all pathways upregulated in that group. Pathway names mentioned in the text are painted red.

In the category of pathways modulated by the virus to support its life cycle, SiPSiC found upregulation of the PI3K/AKT/MTOR and mTORC1 pathways, consistent with evidence that these pathways are activated after SARS-CoV-2 infection (Appelberg et al., 2020). Moreover, mTORC1 activation is known to induce expression of key enzymes involved in several metabolic pathways, among them are glycolysis and biosynthesis of fatty acids and cholesterol (Düvel et al., 2010), supporting SiPSiC’s findings that glycolysis, fatty acid metabolism and cholesterol homeostasis were all upregulated in the alveolar cells of COVID-19 patients. Further support for the upregulation of glycolysis is provided by previous works showing that SARS-CoV-2 infection indeed increases glycolysis, both in colon carcinoma cells and in monocytes (Bojkova et al., 2020; Codo et al., 2020). Importantly, mTORC1 was also shown before to activate the IRE1-JNK signaling pathway of the unfolded protein response (UPR), thereby triggering apoptosis (Kato et al., 2012). SiPSiC found that the UPR pathway was upregulated too (FDR < 1.16 ∗ 10^−68^), consistent with the activation of mTORC1 and previous research suggesting that SARS-CoV-2 induces the unfolded protein response (Echavarría-Consuegra et al., 2021). Interestingly, a previous review stated that UPR activation can increase type 1 interferon production (Sprooten & Garg, 2020). Combined, all these findings suggest that SARS-CoV-2 infection increases endoplasmic reticulum (ER) stress and UPR activation which are also enhanced by the viral-induced upregulation of mTORC1, thereby promoting apoptotic cell death and possibly also the positive feedback loop involving TP53 and type 1 interferons, which further encourages apoptosis. Another pathway in this category which was found upregulated consists of genes upregulated by reactive oxygen species (ROS). Oxidative stress is known to be induced by several viruses (Lee, 2018), and monocytes infected with SARS-CoV-2 showed higher production of mitochondrial ROS (Codo et al., 2020), suggesting that SARS-CoV-2 also induces oxidative stress in the infected cells. Together, these results suggest that SARS-CoV-2 activates MTOR and induces glycolysis and oxidative stress. We applied SiPSiC on human activated B cells (55 cells of COVID-19 patients, 48 control cells) and CD8+ T cells (103 COVID-19 cells, 6 control) to better characterize the adaptive immune response. Despite the relatively small cell populations, TGF beta signaling was found upregulated in activated B cells of COVID-19 patients (FDR < 0.006), consistent with previous reports (Ferreira-Gomes et al., 2021; Melms et al., 2021). The G2/M checkpoint pathway was upregulated in the CD8+ T cell group of COVID-19 patients (FDR < 0.003), a finding missed by the conventional analysis in Melms et al., but consistent with the expected increase in T cell proliferation in the lungs (Liao et al., 2020). The interferon gamma (IFNG) response pathway was found upregulated in the SARS-CoV-2 infected CD8+ T cells as well, albeit with borderline statistical significance (FDR < 0.031). This finding is well correlated with prior evidence of elevated interferon levels in the plasma of COVID-19 patients and the resulting effect on immune cells, and with previous evidence showing upregulation of interferon-related genes across different types of immune cells collected from the mediastinal lymph nodes of SARS-CoV-2 infected monkeys (Schultheiß et al., 2020; Speranza et al., 2021).

Importantly, many of the pathways that SiPSiC found as upregulated in the alveolar cells of the COVID-19 group demonstrated therapeutic potential, including DNA repair, UPR, PI3K/AKT/MTOR signaling, ROS and metabolic pathways. Drugs targeting DNA damage response can block SARS-CoV-2 replication (Garcia et al., 2021), suggesting that SARS-CoV-2 not only increases DNA damage in the infected cells, but also relies on the cells’ reaction to it. SARS-CoV-2 replication can be blocked by UPR inhibitors (Echavarría-Consuegra et al., 2021), PI3K/AKT/MTOR inhibitors (Appelberg et al., 2020; Klann et al., 2020; Stukalov et al., 2021; Yuen et al., 2021), and drugs targeting lipid metabolism (Abu-Farha et al., 2020; Williams et al., 2021). Treatment of COVID-19 patients with N-Acetylcysteine (NAC), a precursor of the antioxidant agent glutathione, was found correlated with lower mortality in a retrospective study (Izquierdo et al., 2022), and SARS-CoV-2 infected monocytes treated with antioxidant agents such as NAC showed reduction in both viral replication and production of several cytokines, in particular interferons of types 1 and 2 (Codo et al., 2020). Codo et al. further showed that the increased production of ROS in infected monocytes promotes glycolysis, and that inhibitors of glycolysis also reduced SARS-CoV-2 replication and could reduce the production of type 1 and 2 interferons. Notably, glycolysis inhibitors also reduced SARS-CoV-2 replication in colon carcinoma cells (Bojkova et al., 2020).

Together this suggests that MTOR signaling and impact on ROS and metabolism play an important role in the pathophysiology of COVID-19, and drugs targeting several of these pathways could have synergistic effects with consequences on disease progression and severity.

In the African green monkey experiment, two monkeys were inoculated with inactivated SARS-CoV-2 (henceforth referred to as control) and eight with the active virus (Speranza et al., 2021). To better study the dynamics of viral infection, these eight monkeys were then split into two groups of four monkeys each and euthanized three (first group) or ten (second) days post inoculation, where the monkeys in the 10 days group had already recovered by the 10^th^ day. Hence, we focused on comparing the active infection (3-days) to the control after calculating the pathway scores per cell type (Methods). Complete results of all three groups can be found in Supplemental Table S2.

Analysis of this dataset provided support for pathways that were found upregulated from all three categories. We found 20 upregulated and 3 downregulated pathways in alveolar cells (Figure 1B). Here too the cells in the active infection group were enriched in the interferon alpha and gamma response, complement, E2F targets and apoptosis pathways relative to both other groups (Supplemental Table S2), supporting our findings from the analysis of human alveolar cells. Importantly, most of the pathways showing therapeutic potential that were found upregulated in the COVID-19 group of alveolar cells in the human dataset were also upregulated in the alveolar cells of the active infection group in the monkey dataset, compared to control. These are the DNA repair, mTORC1 and PI3K/AKT/MTOR signaling, glycolysis, reactive oxygen species and unfolded protein response pathways. Out of all these pathways, only interferon signaling was identified by Speranza et al., further highlighting SiPSiC’s high sensitivity.

Analysis of the immune cells that were found in the monkeys’ lungs revealed that in both the B and CD8+ T cells, the interferon alpha and gamma response pathways were enriched in the 3-days group, in concordance with our finding of interferon gamma upregulation in CD8+ T cells from the human dataset and the evidence showing elevated interferon levels in the plasma of COVID-19 patients and its consequences on immune cells (Schultheiß et al., 2020; Speranza et al., 2021). In addition, both MYC targets pathways (V1 and V2) were found upregulated in the active infection (3 days) group of the CD8+ T cells compared to control. We indeed expect to see upregulation of MYC targets in T cells, since MYC is known to be involved in T cell proliferation (Gnanaprakasam & Wang, 2017), and increased T cell proliferation occurs in COVID-19 patients’ lungs (Liao et al., 2020). To ascertain this finding, we compared *MYC* expression in the control and active infection groups. Indeed, average *MYC* expression 3 days after infection was 89% higher than control (p < 0.0032, unpaired Wilcoxon test), strongly supporting SiPSiC’s finding. Furthermore, the inflammatory response pathway was also enriched in the 3 days group of CD8+ T cells, reflecting the anti-viral activity.

Remarkably, SiPSiC also found the upregulation of the IL6/JAK/STAT3 signaling pathway in the active infection group of CD8+ T cells compared to control (FDR < 5.61 * 10^−9^), a finding supported by evidence for high levels of IL6 in the serum of COVID-19 patients even in relatively mild cases and increased IL6 secretion from epithelial cells infected with SARS-CoV-2 (Han et al., 2020; Patra et al., 2020; Sanli et al., 2021). This is another example for SiPSiC’s potential to highlight therapeutically relevant pathways, as the IL6/JAK/STAT3 signaling pathway has been suggested as a therapeutic target for COVID-19 (Jafarzadeh et al., 2021). Similarly, the Notch signaling pathway was found upregulated in this group, although with borderline statistical significance (FDR < 0.034). Notch too was suggested as a therapeutic target in COVID-19 before, as it is known to modulate the immune response and is involved in a positive feedback loop with *IL6* expression in macrophages (Rizzo et al., 2020). This suggests that CD8+ T cells may exhibit the same feedback loop upon SARS-CoV-2 infection, and that IL6 signaling could be moderated in these cells by inhibiting Notch signaling.

### SiPSiC detects key activated pathways in a tumor specific epithelial lineage of lung adenocarcinoma

To further demonstrate SiPSiC’s robustness, we applied it to lung adenocarcinoma (N. Kim et al., 2020) and compared three malignant epithelial lineages denoted by Kim et al. as tS1, tS2, and tS3 (Supplemental Table S3; 5880 cells total, including 2879 tS1, 2938 tS2, and 63 tS3 cells). While tS1 and tS3 have healthy epithelial counterparts, tS2 showed a tumor specific phenotype, hence we focused on differentially active pathways in this lineage (Supplemental Fig. S1A). In line with the findings of Kim et al., the apoptosis (FDR < 2.4 * 10^−3^) and late response to estrogen (FDR < 3.3 * 10^−4^) pathways were found upregulated in tS2 cells, both compared to tS1 and to tS3. In addition, SiPSiC found upregulation of the hypoxia and glycolysis pathways in tS2 cells (FDR < 1.3 * 10^−6^, FDR < 3.2 * 10^−12^), suggesting hypoxic conditions drive glucose metabolism in tS2 cells. Indeed, a lung adenocarcinoma hypoxia signature (Mo et al., 2020) is upregulated in tS2 cells, supporting our findings. Of note, Mo et al. reported that high expression of this signature was correlated with higher infiltration of activated CD4+ T cells and M0 macrophages, while Kim et al. showed that a positive correlation exists between the proportions of tS2 cells and exhausted CD8+ T cells or monocyte-derived macrophages. Indeed, SiPSiC found upregulation of several inflammatory pathways in the tS2 cells, including the inflammatory response, complement, allograft rejection and IL6/JAK/STAT3 signaling, suggesting an immune response in tS2 cells, agreeing with immune cell infiltration.

Moreover, Kim et al. showed that higher expression of the tS2-specific genes is correlated with metastasis and poorer prognosis. Several pathways SiPSiC identified as upregulated may be involved, including PI3K/AKT/MTOR (FDR < 2.9 * 10^−230^and FDR < 0.019 compared to the tS1 and tS3 lineages, respectively), mTORC1 (FDR < 1.4 * 10^−11^) and epithelial to mesenchymal transition (FDR < 7.7 * 10^−10^), previously demonstrated to be involved in lung adenocarcinoma metastasis (Ding et al., 2018; Krencz et al., 2017; Lu et al., 2020). In addition, cell cycle pathways (G2/M checkpoint, FDR < 3.3 * 10^−4^and E2F targets, FDR < 4.4 * 10^−4^) are upregulated, suggesting tS2 cells proliferate faster and can therefore contribute to tumor aggressiveness.

### SiPSiC analysis of glioma reveals novel differentially active pathways showing potential therapeutic implications

To validate the applicability of SiPSiC to different data types, we analyzed scRNA-seq data of glioblastoma tumors (Neftel et al., 2019). In their work Neftel et al. identified four malignant “meta-modules” (cellular states): Oligodendrocyte-progenitor-like (OPC-like), neural-progenitor-like (NPC-like), astrocyte-like (AC-like) and mesenchymal-like (MES-like). We calculated pathway scores per cell (n = 6,576) for each of the same 50 hallmark pathways. The cells were then split into four groups based on their cell state assignments (1,986 NPC-, 1,047 OPC-, 1,929 AC- and 1,614 MES-like cells), and comparisons were made between each pair of groups (Supplemental Table S4 and Supplemental Fig. S1B). We found that the G2/M checkpoint pathway was upregulated in the OPC- and NPC-like groups compared to the AC- and MES-like groups, indicative of a higher proportion of proliferating cells in these cell states. In addition, the hypoxia response pathway was enriched in the MES-like group compared to all other three groups. These findings are consistent with the findings reported in Neftel et al. Furthermore, SiPSiC analysis suggests that the MES-like group was enriched in the inflammatory response pathway, in concordance with prior evidence that the mesenchymal subtype of GBM tumors is enriched in inflammatory response associated genes (Engler et al., 2012).

Importantly, SiPSiC analysis of the glioblastoma dataset also pointed out pathways with therapeutic implications. The TGF beta (FDR< 4.86*10^−14^), TNFA via NFKB (FDR< 2.77*10^−117^) and IL6/JAK/STAT3 (FDR< 8.86*10^−36^) signaling pathways and the epithelial to mesenchymal transition (EMT) pathway (FDR< 1.02*10^−113^) were all found upregulated in the MES-like group. A previous work has shown that in gliomas, regulatory T cells secrete TGFB1 and thereby promote the NFκB-IL6-STAT3 signaling axis, and that IL6 receptor blockers have a potential therapeutic effect (Liu et al., 2021). TGF beta signaling also activates TNF signaling via NFκB in glioblastoma, which in turn induces mesenchymal transition (Bhat et al., 2013; Yan et al., 2022). Furthermore, these works also suggest that these two pathways can be targeted to improve overall survival and specifically attenuate resistance to radiotherapy in glioblastoma patients. Of relevance, the mesenchymal subtype of glioblastoma is correlated with high levels of both immune markers and infiltration of immune cells (Verhaak et al., 2010; Q. Wang et al., 2017). Together, these findings suggest that regulatory T cells infiltrate glioblastoma tumors, promoting the TGF beta, TNF via NFκB and IL6/JAK/STAT3 signaling pathways and thereby increasing mesenchymal transition of tumor cells.

Furthermore, SiPSiC detected up- and downregulation of the KRAS signaling up (FDR < 9.56 ∗ 10^−22^) and down (FDR < 1.7 ∗ 10^−8^) pathways, respectively, in the MES-Like group, indicative of KRAS signaling activation in it. These findings are consistent with RAS and TGFB1 cooperating to induce EMT in epithelial cells and evidence that KRAS activation in glioblastoma cells induces a mesenchymal shift (H. Kim et al., 2014; Marques et al., 2021; Zhao et al., 2021). SiPSiC also identified upregulation of ROS in the MES-like group compared to all other groups (FDR < 1.05 ∗ 10^−57^), suggesting that the promotion of TGFB1-induced EMT by RAS is mediated by enhanced production of reactive oxygen species, as was shown in mammary epithelial cells (H. Kim et al., 2014). Taken together, our findings suggest a synergistic effect in glioblastoma where regulatory T cells activate TGFB1 signaling leading to EMT which is enhanced by RAS activation. Remarkably, RAS inhibition was shown fatal to glioblastoma cells (Blum et al., 2005). Hence, our analysis suggests that a combined therapy targeting RAS and TGFB1 or their downstream targets could have a synergistic therapeutic effect for glioblastoma, again demonstrating the potential of SiPSiC analysis to accelerate the development of targeted therapy.

Garofano et al. classified the cells into four distinct cell clusters: glycolytic/plurimetabolic (GPM), mitochondrial (MTC), neural (NEU) and proliferative/progenitor (PPR) (Garofano et al., 2021). They found that the GPM and MTC clusters were enriched in cells of the MES- and AC-like cell states, respectively, while the PPR and NEU clusters were enriched in cells of both the OPC- and NPC-like states. Gene set enrichment of these clusters is reported for 40 hallmark pathways, and we compared their results to SiPSiC’s. 31 out of the 40 pathways (78%) were consistent, in the sense that if a pathway was reported as upregulated in one of Garofano et al. cell clusters, SiPSiC also found it is upregulated in the same cell state this cell cluster is enriched in. For instance, Garofano et al. reported that the DNA repair pathway was upregulated in the PPR cell cluster, which is enriched in both OPC- and NPC-like cells. Indeed, SiPSiC found this pathway was upregulated in both the OPC- and NPC-like cell states compared to both the AC- and MES-like states. Furthermore, out of the 9 other pathways eight were found by Garofano et al. to be enriched in the MTC cluster, hence we expected them to be upregulated in the AC-like cell state in our analysis. 7 of these pathways were found to be “near-consistent” in the sense that they were significantly upregulated in the AC-like cell state compared to two of the other three cell states and insignificantly either up- (5 pathways) or down-regulated (2 pathways) in it compared to the third other cell state, making 38 of the 40 (95%) pathways reported by Garofano et al. either completely or near consistent in our SiPSiC analysis.

The two remaining pathways are the hallmark Notch and mTORC1 signaling pathways. Interestingly, while Garofano et al. found that the MTC cluster was enriched in the hallmark Notch signaling pathway (reported FDR < 1.23*10^-3^), SiPSiC analysis found that it was upregulated in the OPC-like cell state (FDR < 0.0095). Although Neftel et al. did not report upregulation of Notch in any of the cell states, AUCell supported SiPSiC’s result finding Notch upregulation in the OPC-like cell state (Supplemental Table S4), but ssGSEA and VAM did not capture this. Importantly, prior research has shown that Notch signaling activation inhibits the maturation and may enhance the proliferation of oligodendrocyte progenitor cells (John et al., 2002; C. Wang et al., 2017; S. Wang et al., 1998), suggesting that Notch upregulation in OPC-like glioblastoma cells may play similar roles in the pathophysiology of the disease. Similarly, while Garofano et al. found that the mTORC1 signaling pathway was enriched in the PPR cluster (reported FDR < 1.26*10^-3^) and Neftel et al. did not report its upregulation in any of the cell states, SiPSiC found that it was upregulated in the MES-like cell state (FDR < 3.1*10^-75^), supported by AUCell and VAM which also found the same (Supplemental Table S4). Prior research has found that in epithelial cells, TGFB1-induced EMT both promotes and requires the activation of the PI3K/AKT/MTOR pathway, and more specifically activates the mTORC1 complex (Lamouille et al., 2014). Importantly, the same has also been suggested for an EMT-like process in glioblastoma (Iser et al., 2017; L. Zhang et al., 2014). Intriguingly, SiPSiC found the upregulation of the PI3K/AKT/MTOR signaling pathway in the MES-like cell state (supported by AUCell and ssGSEA), in addition to the upregulation of the TGF beta signaling and EMT which were mentioned above (both supported by AUCell, ssGSEA and VAM, see Supplemental Table S4). Combined, these findings may well account for upregulation of the mTORC1 signaling in this cell state, supporting our finding over what Garofano et al. reported for this pathway.

As further validation, we applied SiPSiC to lower-grade oligodendroglioma scRNA-seq dataset (Tirosh et al., 2016). We compared SiPSiC scores of three subpopulations: 924 cancer stem cells (CSC), 1300 astrocyte-like cells and 1820 oligodendrocyte-like cells (Supplemental Table S5 and Supplemental Fig. S1C). SiPSiC identified upregulation of the G2/M checkpoint pathway in the CSC group (FDR < 3.3 ∗ 10^−12^), consistent with the findings of Tirosh et al. Furthermore, SiPSiC identified other relevant differential pathways not reported in the original analysis. The Wnt/beta catenin signaling (FDR < 1.6 ∗ 10^−6^) and DNA repair pathways (FDR < 1.3 ∗ 10^−6^) were upregulated in the CSCs compared to the two other groups, consistent with evidence that Wnt signaling plays a central role in the maintenance of stem cells in several tissues and the stemness of glioma cells in particular (Fodde & Brabletz, 2007; Jin et al., 2011; Zheng et al., 2010) and that in glioma stem cells, activation of the DNA damage response confers radioresistance (Bao et al., 2006). Additionally, the CSC subpopulation showed upregulation of the E2F targets (FDR < 1.5 ∗ 10^−26^) and MYC targets pathways (FDR < 4.9 ∗ 10^−12^ for MYC targets V1 and FDR < 0.0009 for MYC targets V2). *MYC* is more highly expressed in glioma CSCs compared to non-stem glioma cells, and knockdown of *MYC* reduces CSC proliferation and promotes apoptosis (J. Wang et al., 2008). Lymphoid-specific helicase (HELLS) activity is essential for glioblastoma stem cells and correlates with MYC and E2F targets (G. Zhang et al., 2019), suggesting SiPSiC captures elevated HELLS activity in CSCs.

Interestingly, SiPSiC analysis suggested that the astrocyte-like cells in oligodendroglioma also harbor a mesenchymal phenotype. 20 out of 23 pathways (87%) that were upregulated in the MES-like glioblastoma cells mentioned above were also upregulated in the astrocyte-like cells of oligodendroglioma, including EMT, hypoxia, glycolysis, and several inflammation-related pathways, a significant overlap (p < 0.005, Fisher exact test). Similarly, all 4 pathways upregulated in the AC-like glioblastoma cells were also upregulated in the astrocyte-like cells of oligodendroglioma, providing further evidence of their astrocytic nature.

To conclude, SiPSiC differential pathway analyses of the COVID-19 and glioma datasets demonstrate that SiPSiC can accurately detect key differentially active pathways even in sparse datasets with few cells or when many of the pathways’ genes are not available in the data. This allows for comprehensive and sensitive pathway analysis in single cells, that can serve to study fundamental biology as well as generate clinically and therapeutically relevant hypotheses, that may be missed by standard analysis.

### Unsupervised clustering by SiPSiC scores identifies shared biological features and cellular identities

Unsupervised clustering of scRNA-seq datasets allows to demarcate distinct cell subpopulations within tissues, offering insights into tissue heterogeneity in various scenarios. In order to assess the potential benefits of clustering based on SiPSiC pathway scores compared to the conventional approach of clustering by gene expression, we calculated cell clusters and Uniform Manifold Approximation and Projection (UMAP) for 9635 malignant glioblastoma cells (Neftel et al., 2019), both by gene expression and by SiPSiC scores (Methods).

Gene-based clustering produced 13 clusters. Although the dataset was sampled from 9 different patients, in 5 clusters (0, 1, 5, 9 and 10) more than 99% of the cells were of a single patient, and in 1 other (cluster 7) 97% of the cells were of a single patient, reflecting the strong patient-bias typical for gene-based clustering of malignant cells (Figure 2, panels A, B). In contrast, using SiPSiC pathway scores for clustering with the same algorithm and resolution produced 6 clusters. In 5 of these clusters no more than 76% of the cells were of a single patient, suggesting that patient specific batch effects have limited effect on SiPSiC based clustering (Figure 2, panels A-B). While 96% of the cells in cluster 0 belong to patient 105, most of these cells are classified as MES-like1 cells, suggesting this cluster may capture their MES-like1 identity and not necessarily patient specific artifacts. Together, this suggests that SiPSiC based clustering was largely based on similar biologically relevant features of the cells rather than patient specific batch effects.

**Figure 2.**
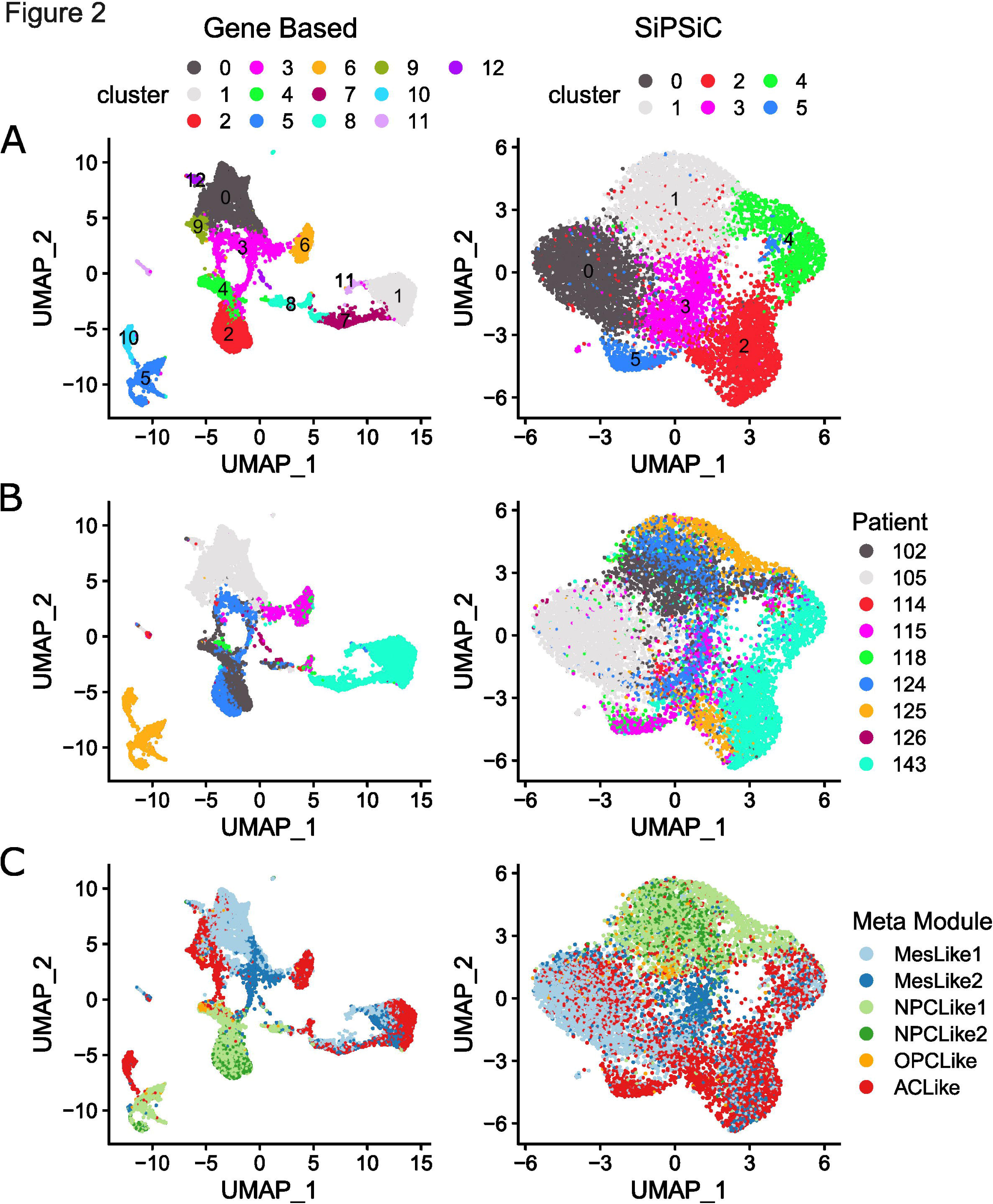
**(A-C)** UMAP projections based on gene expression (left) and hallmark pathway scores by SiPSiC (right). **(A)** Cells were clustered by Louvain algorithm either according to SiPSiC scores or to gene expression. UMAPs show cells colored by cluster. **(B)** Cells colored by patient identity. **(C)** Cells colored by malignant meta-module assignment.

This also allows to detect functional heterogeneity in an unsupervised manner by comparing differential pathway scores between the clusters (see Supplemental Table S6). Cluster 1 was enriched in the Wnt/beta catenin and hedgehog signaling pathways, both found upregulated in the NPC-like meta-module in our differential pathway analysis of the Smart-seq2 glioblastoma dataset above (Supplemental Fig. S1B). Indeed, 2096 (84%) out of the 2496 cells in this cluster were NPC-like cells (Figure 2C). Furthermore, the adjacent cluster 4 contains 339 additional NPC-like cells, forming a distinct group of 2435 (96.4%) of the 2527 NPC-like cells in the data on the UMAP projection. Similarly, 12 out of the 13 pathways found enriched in cluster 3 were detected by SiPSiC as upregulated in the MES-like cells, and 883 (76%) out of the 1164 cells in this cluster are MES-like cells. Importantly, the adjacent clusters 0 and 2 contain 1977 and 644 additional MES-like cells, respectively, forming together with cluster 3 a large group of 3504 (87%) of the total 4017 MES-like cells on the UMAP projection (Figure 2C). Moreover, 3 out of the 4 pathways that were enriched in cluster 5 showed upregulation in AC-like cells: The interferon alpha and angiogenesis pathways are significantly upregulated in the AC-like cells vs. all other meta-modules, and the interferon gamma pathway that is significantly upregulated compared to NPC- and OPC-like cells (Supplemental Fig. S1B and Supplemental Table S4). Indeed, 356 (78%) of the 455 cells in this cluster are AC-like cells (Figure 2C).

In contrast, testing the gene-based clusters for enrichment of these cellular identities, we found that both the MES-like and NPC-like cells were split across different groups of clusters. While clusters 0, 3 and 9 contain 2754 (69%) of the MES-like cells in the data, additional 955 (24%) MES-like cells are found in clusters 1, 7 and 11 (Figure 2 panels A, C). Of note, less than 21% of the cells in cluster 9 are MES-like cells. NPC-like glioblastoma cells are even more scattered across different clusters. While 1680 (66%) of the NPC-like cells are found in clusters 2 and 4, clusters 5 and 8 contain additional 570 (23%) and 170 (7%) NPC-like cells, respectively, resulting in 3 clearly distinct groups of NPC-like cells on the UMAP (Figure 2 panels A, C). 99.8% of the cells in cluster 5 belong to a single patient, again showing strong patient bias that hinders the ability to cluster the cells properly.

Together, these results demonstrate that SiPSiC scores emphasize common cellular functions and are robust to batch effects, and are therefore better fitted to uncover the biological underpinnings of different cell clusters. Other methods were developed to explicitly remove known batch effects such as patient specific differences. To compare SiPSiC clustering results to explicit patient batch correction we applied Seurat integration (Butler et al., 2018), Harmony (Korsunsky et al., 2019) and scVI (Lopez et al., 2018) to the same dataset and repeated the clustering pipeline (Supplemental Fig. S2). All three methods removed the differences between patients, similarly revealing the functional clustering discovered by SiPSiC. Therefore, while we do not suggest SiPSiC as a batch correcting tool per se, we do point to its utility in focusing on biologically relevant cellular characteristics. Moreover, it can be useful even in the context of overcoming artifacts, especially when the batches and the source of the artifacts are unknown and cannot be directly removed, or when batch correction over-corrects real biological differences.

### Pathway scoring methods benchmarking reveals higher accuracy and improved scalability of SiPSiC for different data types

To compare SiPSiC results to state of the art methods for scoring pathway activity in single cells, we repeated the differential pathway analyses of the glioblastoma and both COVID datasets with AUCell (Aibar et al., 2017), Variance-Adjusted Mahalanobis (VAM) (Frost, 2020) and ssGSEA (Barbie et al., 2009).

Summary of differentially active pathways between the actively infected monkeys and controls by all four methods can be found in Table 1. All four methods found upregulation of the mTORC1 signaling, MYC targets V1, oxidative phosphorylation, ROS, allograft rejection and both interferon pathways in the active infection group of the alveolar cells. However, several differences were observed between the different methods. SiPSiC and ssGSEA were the only methods to detect significant upregulation of apoptosis (Figure 3), while SiPSiC was the only method to detect upregulation of the adipogenesis pathway in alveolar cells. Both findings are supported by all four methods detecting the same in the active infection group of the human alveolar cells.

**Figure 3.**
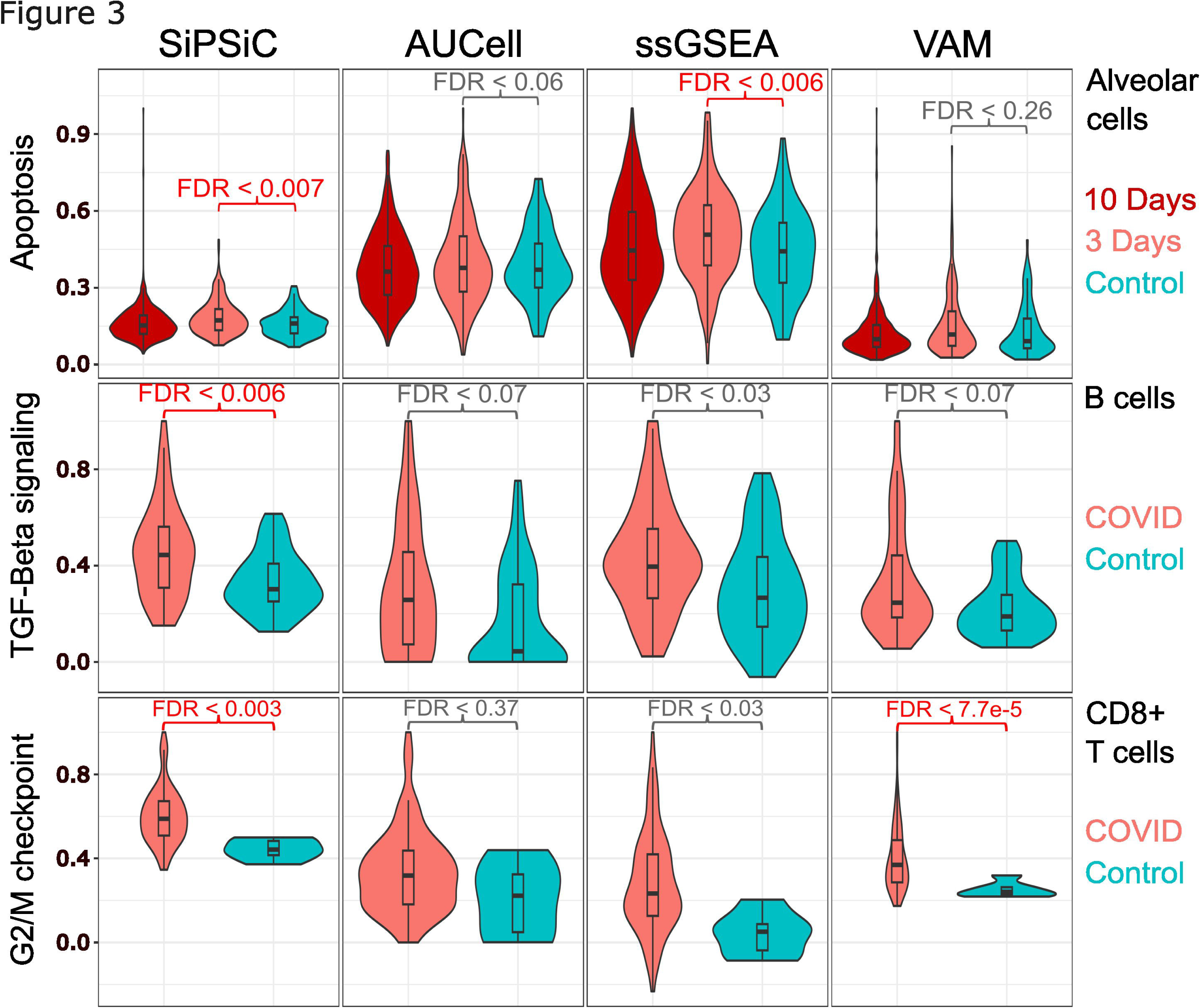
Violin plots showing normalized pathway scores distribution as calculated by the four different methods. Significant results (FDR < 0.01) are painted red. Upper panel – Apoptosis, monkey alveolar cells. Middle – TGF beta signaling, human B cells. Lower – G2/M checkpoint, human CD8+ T cells.

**Table 1.** Summary of the number of differentially active pathways in the active infection group of the COVID monkey dataset compared to control, by all four tested methods. The overlap column shows the number of pathways found as differentially active by all methods. Skipped pathways are pathways for which AUCell could not calculate scores, while missed pathways are cases where the relevant method failed to detect differential active of a pathway detected by all other methods. See Supplemental Table S2 for the analysis of all 50 hallmark pathways.

| Cell Type | Pathway Status | SiPSiC | AUCell | ssGSEA | VAM | Overlap |
| --- | --- | --- | --- | --- | --- | --- |
| Alveolar | Upregulated | 20 | 9 | 18 | 11 | 7 |
|  | Downregulated | 3 | 10 | 9 | 0 | 0 |
|  | Skipped | 0 | 2 | 0 | 0 | 0 |
|  | Missed | 0 | 3 | 1 | 2 | - |
| B Cells | Upregulated | 2 | 0 | 4 | 2 | 0 |
|  | Downregulated | 7 | 0 | 6 | 2 | 0 |
|  | Skipped | 0 | 20 | 0 | 0 | 0 |
|  | Missed | 0 | 4 | 0 | 0 | - |
| T Cells | Upregulated | 20 | 12 | 17 | 11 | 7 |
|  | Downregulated | 4 | 4 | 6 | 0 | 0 |
|  | Skipped | 0 | 16 | 0 | 0 | 0 |
|  | Missed | 0 | 3 | 0 | 2 | - |

To compare the methods in an unbiased manner, we defined misses of a method as pathways whose differential activity was consistently detected by all three other methods (FDR < 0.01), but not by the method in question (either did not pass FDR or changed in the opposite direction). For instance, AUCell failed to detect differential activity of the complement pathway (FDR < 0.5) in the active infection group of the monkey alveolar cells, whereas all other methods showed significant upregulation. AUCell also failed to detect differential activity of the mitotic spindle, interferon alpha and gamma pathways in B cells, while all other methods found upregulation of the interferon pathways and downregulation of the mitotic spindle pathway, suggesting SARS-CoV-2 infection hinders B cell proliferation in the host’s lungs.

Comparing the results of the CD8+ T cell analyses, we found that all methods successfully detected the activation of cells, as all four showed upregulation of the TNFA via NFKB, complement, inflammatory response, and interferon pathways in this group. Importantly, SiPSiC was the only method to detect upregulation of both MYC targets pathways in the active infection group, while AUCell missed the MYC targets V2 pathway, producing a median score of zero for all T cell groups, and the two other methods failed to reach statistical significance for the MYC targets V1 pathway (Supplemental Fig. S3). As mentioned above, CD8+ T cell proliferation can be expected in the lungs of COVID-19 patients, and a significantly higher (89%) average *MYC* expression was indeed found in the active infection group compared to control in this dataset (p < 0.0032, unpaired Wilcoxon test), supporting SiPSiC’s higher robustness.

For the human COVID dataset we found the methods largely agree on the upregulated pathways in alveolar cells of COVID-19 patients, with 24 pathways detected by all four (Table 2). The inflammatory response, IL6/JAK/STAT3 signaling and TNFA signaling via NFKB were all downregulated according to all four methods, suggesting a counterintuitive reaction of infected epithelial cells suppressing immune response. Additionally, the TGF beta signaling pathway was found downregulated by all methods except ssGSEA, providing further support for this putative anti-inflammatory response. In activated B cells, SiPSiC was the only method that managed to show upregulation of TGF beta signaling (Figure 3), consistent with previous reports as mentioned above, whereas all other methods did not find any differentially active pathway in this group. Similarly, SiPSiC and VAM were the only methods to detect upregulation of the G2/M checkpoint pathway in CD8+ T cells (Figure 3), in line with the upregulation of *MYC* found in the monkey T cells and the expected proliferation of these cells as mentioned above.

**Table 2.** Summary of the number of differentially active pathways in COVID patients compared to control, by all four methods. The overlap column shows the number of pathways found as differentially active by all methods. Skipped pathways are pathways for which AUCell could not calculate scores, while missed pathways are cases where the relevant method failed to detect differential activity of a pathway detected by all other methods. See Supplemental Table S1 for the analysis of all 50 hallmark pathways.

| Cell Type | Pathway Status | SiPSiC | AUCell | ssGSEA | VAM | Overlap |
| --- | --- | --- | --- | --- | --- | --- |
| Alveolar | Upregulated | 31 | 30 | 34 | 28 | 24 |
|  | Downregulated | 12 | 9 | 8 | 10 | 4 |
|  | Skipped | 0 | 4 | 0 | 0 | 0 |
|  | Missed | 0 | 4 | 1 | 3 | - |
| B Cells | Upregulated | 1 | 0 | 0 | 0 | 0 |
|  | Downregulated | 0 | 0 | 0 | 0 | 0 |
|  | Skipped | 0 | 19 | 0 | 0 | 0 |
|  | Missed | 0 | 0 | 0 | 0 | - |
| T Cells | Upregulated | 1 | 0 | 1 | 3 | 0 |
|  | Downregulated | 0 | 0 | 0 | 0 | 0 |
|  | Skipped | 0 | 24 | 0 | 0 | 0 |
|  | Missed | 0 | 0 | 0 | 0 | - |

For the glioblastoma data, we found less agreement between the methods (see Table 3). SiPSiC showed no misses, demonstrating its higher robustness across data types, however AUCell missed 3 pathways while ssGSEA and VAM missed 5 pathways each, where most misses are due to biases towards specific meta-modules. VAM showed bias towards the MES-like group (28 upregulated pathways compared to 20, 23 and 17 detected by AUCell, SiPSiC and ssGSEA, respectively) and NPC-like group (13 upregulated pathways compared to 6, 8 and 4 detected by AUCell, SiPSiC and ssGSEA, respectively). Specifically, VAM detected upregulation of the adipogenesis pathway in the MES-like group, while all other methods found it was upregulated in the AC-like group. This bias prevented VAM from detecting differential pathway activity in either the MES- or NPC-like group over the other, which accounts for 3 more misses of the method. Indeed, while all other methods found upregulation of the PI3K/AKT/MTOR signaling and heme metabolism pathways in the MES-like group, and upregulation of the hedgehog signaling in the NPC-like group, VAM detected upregulation of the PI3K/AKT/MTOR signaling pathway in the NPC-like group, while it failed to achieve statistical significance comparing the NPC- and MES-like groups in the two other pathways. In summary, these biases account for 4 out of the 5 misses of VAM.

**Table 3.** Summary of the number of differentially active pathways in each glioblastoma meta-module, according to the four tested methods. Missed pathways are cases where the relevant method failed to detect differential activity of a given pathway in the specific cell group where all other methods detected the pathway. See Supplemental Table S4 for a summary of the analysis of all 50 hallmark pathways.

|  | SiPSiC | AUCell | ssGSEA | VAM | Overlap |
| --- | --- | --- | --- | --- | --- |
| AC-like | 4 | 13 | 10 | 1 | 1 |
| MES-like | 23 | 20 | 17 | 28 | 13 |
| OPC-like | 2 | 2 | 10 | 0 | 0 |
| NPC-like | 8 | 6 | 4 | 13 | 2 |
| Missed | 0 | 3 | 5 | 5 | - |

ssGSEA, in contrast, showed bias towards the OPC-like group with 10 pathways detected as upregulated compared to 2, 2 and 0 detected by AUCell, SiPSiC and VAM, respectively. This bias accounts for 4 out of ssGSEA’s 5 misses, as it found upregulation of the Wnt/beta catenin, G2/M checkpoint, and E2F targets pathways in the OPC-like group, all found upregulated in the NPC-like group by the other methods, and failed to show significant difference between the OPC- and MES-like groups in the mTORC1 signaling pathway which was found upregulated in the MES-like group by all other methods. A similar bias was presented by AUCell towards the AC-like group with 13 pathways the method found as upregulated in it compared to 4, 10 and 1 found by SiPSiC, ssGSEA and VAM, respectively. This bias accounts for 2 of AUCell’s 3 misses, one of them being the inflammatory response pathway which all other methods detected as upregulated in the MES-like group, consistent with previous knowledge as mentioned above.

Strikingly, SiPSiC was the only method with no misses across all cell types in all three datasets, suggesting it is the most robust method among the four (see Tables 1 through 3).

We also compared the execution time of the different methods, applied to the same three datasets (see Methods). The detailed execution times and fold change figures compared to SiPSiC can be found in Supplemental Table S7. AUCell, ssGSEA and VAM were at least 43%, 169% and 77% slower than SiPSiC across all cell types and datasets, respectively, excluding the B cells of monkeys where AUCell was slightly quicker than SiPSiC. On average, we saw a fold change of 1.99, 11.06 and 3.71 in the execution times of AUCell, ssGSEA and VAM across the datasets, compared to SiPSiC execution times. Importantly, the differences in execution times were largely correlated with dataset size so that SiPSiC’s advantage was greater in larger datasets. For instance, while AUCell, ssGSEA and VAM showed a fold change of 0.77, 2.69 and 1.77 compared to SiPSiC for the B cells of monkeys (113 cells), these numbers increased to 1.85, 9.25 and 2.33, respectively, in the analysis of monkey alveolar cells where 1004 cells were included. Moreover, in the glioblastoma analysis (6576 cells) SiPSiC executed in just 5.7 seconds, whereas AUCell, ssGSEA and VAM took 14.1 (247% of SiPSiC’s execution time), 146.2 (2565%) and 20.5 (360%) seconds to complete. Therefore, SiPSiC is more scalable and better fitted for use on large datasets, allowing meta-analysis of large amounts of single cell data becoming available now and in the near future.

### SiPSiC is robust to normalization parameter settings

SiPSiC normalizes the expression of each gene by the median expression level of the top τ% of cells (see Methods). The above analysis was conducted with the default value of τ=5%. To test the robustness of SiPSiC results to τ we repeated the analyses of the same 5 datasets with τ=2% and τ=10% (Supplemental Table S8). The different values of τ had only minimal impact on the results and most pathways were either found not significantly different, or significantly upregulated in the same group of cells. In the lung adenocarcinoma and both COVID datasets, between 46 (92%) and 49 (98%) of the 50 hallmark pathways produced the same results across all comparisons. For glioblastoma and oligodendroglioma 38 (76%) and 43 (86%) of the pathways produced the same result.

Across the datasets, we found only 3 pathways where results with τ=5% were inconsistent with both τ=2% and τ=10%. These are the interferon gamma pathway in the glioblastoma analysis, the allograft rejection in the T cells of monkeys and the genes downregulated by UV in T cells of human SARS-CoV-2 patients. In glioblastoma upregulation of interferon gamma response in AC-like cells was observed with τ=2%, and in MES-like cells with τ=10%, while with τ=5% both groups were found upregulated compared to OPC-like and NPC-like cells, as part of an overall bias towards AC-like cells upregulation with τ=2% and MES-like cells with τ=10% (Supplemental Table S8). For the allograft rejection pathway changes were minimal, but just crossed the selected threshold of FDR < 0.01 (0.0094 for τ=2%, 0.0112 for τ=5%, and 0.0089 for τ=10%). For genes downregulated by UV, downregulation in T cells was detected with τ=2% and τ=10%, but was not significant with τ=5% as well as according to AUCell, ssGSEA and VAM, suggesting the results with τ=5% better reflect reality.

## Discussion

scRNA-seq is a powerful technique to interrogate cellular heterogeneity, allowing researchers to comprehensively query changes in biological processes in high resolution. Despite its utility, computational methods for inferring the activity of biological processes are still lacking. As the scale, resolution and abundance of scRNA-seq continue to grow, there is a pressing need to refine existing methodologies and develop novel improved ones. These endeavors are pivotal for uncovering novel biological processes and elucidating poorly understood phenomena.

In this paper we introduce SiPSiC, a novel method for inferring pathway scores from scRNA-seq data. We demonstrate its applicability and high sensitivity, accuracy, and consistency by applying it to publicly available datasets of COVID-19 and various malignancies. Our analyses reveal numerous pathways exhibiting altered activity, many of which were not previously identified in the original papers. Importantly, these findings align with established biological knowledge and are corroborated by other research studies, underscoring the biological relevance of SiPSiC-derived insights.

SiPSiC presents several advantages over the conventional approach of gene set enrichment of differential genes. Its primary benefit lies in its capability to assess pathway activity for every single cell, allowing more robust detection of changes in pathway activity, estimation of heterogeneity in pathway activity changes and representation of the data in pathway space. We suggest a novel approach for dimensional reduction and clustering for single cell data using SiPSiC scores rather than gene expression profiles. This strategy accentuates biological similarities between cells, while mitigating technical artifacts and covariates such as patient of origin. While SiPSiC is not a batch correction method, it offers distinct advantages that may render batch correction unnecessary in certain scenarios. Unlike batch correction methods, SiPSiC does not artificially alter the raw data, thus avoiding inadvertent removal of genuine biological differences. Moreover, batch correction requires knowing what the batches are, and that they are at least partially independent of the biological differences, and therefore may not always be feasible.

Comparative analyses against existing methods that can compute pathway activity per cell-AUCell, ssGSEA and VAM, demonstrate SiPSiC’s superior ability to identify real, biologically meaningful results, while substantially reducing computational execution time. In summary, we demonstrate that SiPSiC provides accurate, comprehensive, and useful insights into biological pathway activity at the single cell level. We attribute this success to the combination of proper normalization ensuring comparable contribution from different genes, and rank-based weighting allowing to take advantage of the higher information content of highly expressed genes.

However, our algorithm does present potential pitfalls. First, SiPSiC’s results depend on the units in which gene expression data are provided. Based on our analyses, we recommend providing SiPSiC with TPM values (or CPM if normalization by gene length is not required) rather than logarithmic transformations of these values. Second, SiPSiC is susceptible to outliers, and we therefore recommend to filter both genes and cells before applying it. Lastly, the activation of many biological processes involves the induction of some genes and the silencing of others. SiPSiC does not account for genes changing in the opposite direction in the same gene set, hence gene sets used as input for SiPSiC analysis should be separated according to the direction of change upon activation of the pathway. To address this, we selected the MSigDB hallmark pathway database, whose pathways meet this criterion (Liberzon et al., 2015). While other methods developed for bulk RNA-seq data can model more complex pathway structures (Drier et al., 2013; Tarca et al., 2009; Vaske et al., 2010; Young & Craft, 2016), they rely on high quality data and therefore are prone to errors when applied on sparse and noisy data typically achieved by single cell RNA-seq.

## Methods

### The SiPSiC algorithm

Taking an scRNA-seq gene expression matrix *X* in TPM or CPM, and a given gene-set, SiPSiC performs the following steps to calculate the score for all the cells in the data:

(1) **Calculate normalized gene scores**: Calculate the median of the τ percent of cells with highest expression (default τ=5%). If it is positive, use it as normalization factor, if zero use the maximum value as the normalization factor. Calculate new normalized gene scores for each gene *i* in cell *j:* 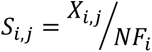 where *NF_i_* is the normalization factor.

The reason behind this step is that scRNA-seq data are normally sparse (Chi & Deng, 2020), namely, the fraction of zeros in the data is large. Thus, by selecting the median of the top τ% cells there is a high likelihood that for most genes the value will be greater than zero, while on the other hand it will also not be an outlier, which may perturb further processing steps.

(2) **Pathway scoring**: Rank the genes in the gene-set by their total expression across all cells ∑_*j*_ *X_i,j_*. The pathway score Pj is the weighted average of the normalized gene scores by the 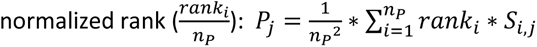.

### scRNA-seq datasets preprocessing

We downloaded published datasets from five papers (N. Kim et al., 2020; Melms et al., 2021; Neftel et al., 2019; Speranza et al., 2021; Tirosh et al., 2016). For the Human COVID, lung adenocarcinoma and oligodendroglioma datasets, cell identity annotations were downloaded as well. For the glioblastoma Smart-seq2 dataset we used the malignant cell meta-module assignment that was published. To identify cell identities for the monkey COVID-19 dataset, we first identified biomarkers for alveolar, activated B, and CD8+ T cells. First, we used the human COVID-19 dataset to extract marker genes of each cell type compared to all other cells, using the FindMarkers function of Seurat (Butler et al., 2018), with logfc.threshold = 1 and min.diff.pct = 0.5, following the parameters used by Speranza et al. In addition to these markers we also selected canonical marker genes of alveolar, B and T cells from the literature: CAV1, PDPN (*T*1*α*), SFTPB, SFTPC and SFTPD were used for alveolar cells (Chen et al., 2004), CD20, CD27, CD28 for B cells (Sanz et al., 2019) and CD2, CD8 and CXCR3 (CD138) for T cells (Groom & Luster, 2011; Tzankov et al., 2005). Next, we clustered the monkey dataset using R’s Seurat clustering pipeline described below with a 0.5 resolution (for the findClusters function), and all clusters were annotated as a given cell type if and only if they displayed high expression of both the canonical markers and the markers identified in the human dataset. This resulted in 1411, 1060 and 17365 cells identified as alveolar, B, and CD8+ T cells, respectively, before filtering was applied. After filtering, these numbers dropped to 1004 (86 control, 217 3-days, and 701 10-days) alveolar cells, 113 (29 control, 71 3-days, and 13 10-days) B cells, and 2224 (360 control, 1162 3-days, and 702 10-days) T cells.

The glioblastoma Smart-seq2, lung adenocarcinoma and oligodendroglioma datasets were log normalized by the original authors, hence we first converted them to linear scale. For the glioblastoma 10x Genomics dataset, we used all malignant cells and genes reported by Neftel et al. after their filtering. For all other datasets, we took all relevant cells (see below), removed cells with less than 1,000 expressed (at least 1 read) genes, and then excluded genes expressed in less than 10% of the remaining cells. For the two COVID-19 datasets, we selected Alveolar, B cells and CD8+ T cells, and applied above filtering for each cell type separately. Since both datasets were supplied as simple count matrices, we divided all transcripts by the total counts of their cell of origin and multiplied by 1 million after filtering. For the lung adenocarcinoma dataset, we selected only tS1, tS2 and tS3 cells. For oligodendroglioma we selected only malignant cells.

### SiPSiC score comparisons

We applied SiPSiC for each dataset separately. Unpaired Student’s t-test was used to compare SiPSiC scores between two groups of cells. When more than two groups of cells were identified in the dataset (lung adenocarcinoma, glioma, and…) we performed all pairwise comparisons. The reported FDR values represent the largest FDR of all pairwise comparisons. We considered a pathway upregulated (downregulated) only when there was a positive (negative) median difference and an FDR < 0.01 compared to each of the other groups, unless stated otherwise.

To infer the Mo et al. hypoxia signature was upregulated in the tS2 lineage, we relied on the fact that all four genes included in this signature were tS2-specifc (supplementary data 3, Kim et al.). In the glioblastoma dataset analysis, since Garofano et al. performed pathway analysis on different groups of cells than the ones reported in Neftel et al. we relied on the overlaps between the two classifications. Since the PPR and NEU clusters were enriched in both NPC- and OPC-like cells, when a specific pathway was reported as upregulated in one of these clusters, we considered our result consistent with the findings of Garofano et al. even when only one of the respective cell states (either OPC- or NPC-like) showed upregulation of that pathway compared to the MES- and AC-like cell states. Such “semi consistent” pathways account for 5 of the 31 consistent pathways mentioned in the results section.

Heatmaps were generated using the R package pheatmap, version 1.0.12. Only differential pathways are shown. After selecting relevant cell groups (see below), we calculated z-scores from SiPSiC pathway scores for each of the pathways individually. We sorted the cells in each cell group based on their average z-score across all pathways upregulated in that group and plotted them in ascending order (left to right). The pathways upregulated in each group were sorted in ascending order of FDR values (most significant result at the top). Following the emphasis in the main text, only the active infection (3 days) and control groups were selected for the monkey COVID dataset, while for the lung adenocarcinoma dataset we only compared the tS1 and tS2 cell lineages, as the tS3 lineage contained only 63 cells. Since almost all pathways showed a significant difference between these two groups, we included the 20 most significant pathways in the heatmap, and sorted the cells in each of the two lineages according to their average z-scores across these 20 pathways.

### Clustering and cluster composition analysis

We started with all gene expression data provided in the glioblastoma 10X Genomics dataset published by Neftel et al. To distinguish between malignant and non-malignant cells, we first inferred malignant cell markers from the Smart-seq2 dataset according to the cell annotation provided by Neftel et al. Markers were calculated using the Seurat FindMarkers function with all parameters set to default, and the 10 genes with the highest fold change that passed the adjusted p < 0.01 threshold were selected as the final malignant markers. This resulted in the following genes: PTPRZ1, IGF2, FABP7, GPM6A, CHI3L1, EGFR, IGFBP3, BCAN, NNAT and PDGFRA. We then calculated for each cell in the 10x Genomics dataset the average expression (in log(TPM +1) units) of these 10 markers as well as the markers of T cells, macrophages and non-malignant oligodendrocytes provided by Neftel et al. We clustered all cells using the Louvain algorithm by applying the Seurat FindClusters function with a 0.5 resolution using the first 15 principal components. This yielded 12 clusters, each showing either upregulation of malignant markers or markers of one the non-malignant cell types. For downstream analysis we focused on the 9635 cells in the clusters marked as malignant.

These cells were then clustered twice using Seurat (version 4.4.0) with identical parameters, first based on gene expression and second based on SiPSiC’s pathway scores. The first 15 principal components were identified using RunPCA, and used to find the 20-nearest neighbors using FindNeighbors (with dims = 15). Cells were clustered by the Louvain algorithm as implemented in the FindClusters function with a resolution of 0.3. Differential pathways were identified in the pathway-based clustering, using the Wilcoxon rank sum test implemented in Seurat’s FindAllMarkers function, using parameters slot = counts, only.pos = true and logfc.threshold = 0.03. All UMAP projections were calculated by RunUMAP based on the first 15 principal components.

We used the marker genes provided by Neftel et al. for each malignant meta-module to annotate the cells, using the Seurat addModuleScore function with ctrl = 50, assigning each cell to the meta-module with the maximal score. This annotation resulted in 2919 AC-like, 2000 NPC-like1, 527 NPC-like2, 2869 MES-like1, 1148 MES-like2 and 172 OPC-like cells. Throughout the clustering section we used the terms NPC- or MES-like cells to refer to the combination NPC-like1 and NPC-like2 cells or MES-like1 and MES-like2 cells respectively.

To integrate the data for batch correction using Seurat we implemented the standard Seurat integration pipeline. We started by finding the 2000 most variable genes in each patient’s normalized data using the SplitObject, NormalizeData and FindVariableFeatures functions, then detected the relevant features for integration using the SelectIntegrationFeatures and FindIntegrationAnchors functions with default values. Lastly, we integrated the data by executing the IntegrateData function with k.weight = 70. We then clustered the cells using the same functions and parameters described above for the clustering based on SiPSiC scores.

To apply the scVI integration method, we followed the scVI documentation and used the raw counts of the same 10x Genomics dataset rather than the TPM values we used in all other integration methods. Integration was applied using the python package scVI-tools, version 1.1.2, and python version 3.9.18. We first selected the 2000 most variable genes using the FindVariableFeatures function, then followed the standard scVI pipeline by calling the setup_anndata function with batch_key set to the patient identity as well as the scVI and train functions to create and train the model required for integration. We proceeded with calling the get_latent_representation function then the Seurat CreateDimReducObject, and finally executed the FindNeighbors and RunUMAP functions with dims = 1:10 and the scVI reduction as input for the reduction parameter, as instructed by the scVI documentation. For the Harmony integration we used the R harmony package version 1.2.0. We executed the RunHarmony function with the patient identity variable name as input and all other values set to default. We then clustered the cells using the same functions and parameters described above for the clustering based on SiPSiC scores.

### Method benchmarking

We repeated the SiPSiC analyses for the glioblastoma (Smart-seq2) and two COVID datasets using AUCell version 1.24.0 and VAM 1.1.0. ssGSEA was applied using its implementation in R’s GSVA package (Hänzelmann et al., 2013), version 1.50.0. To guarantee integrity of the results, the same preprocessing steps were applied, cell assignments to the different groups were kept and pathway up- or down-regulation were defined as in our SiPSiC analyses across the datasets. All parameters of the different methods were set to default. Pathways skipped by AUCell but consistently detected by all other methods as either up- or downregulated in a specific group were counted both as skipped and missed by AUCell.

Violin plots were generated with the R package ggplot2, version 3.4.4. For each method, all cell scores were normalized with (divided by) the maximum score produced by this method for the relevant pathway prior to plot generation.

We used R studio’s profiler (R’s Profvis package, version 0.3.8) to record the execution of the different methods on a computer with windows 11 x64, 6 intel i7-8700 (3.20 GHz) cores and 64 GB RAM. The R version was 4.3.2 used in R studio version 2023.12.0 (build 369). SiPSiC execution times were calculated as the sum of all calls to the getPathwayScores function for each of the 50 hallmark pathways, while for AUCell we measured the execution time of the AUCell_run function. Since these functions also include the detection of the relevant pathway genes in the input data matrix, for VAM we summed the executions of the createGeneSetCollection and vamForCollection functions, while for ssGSEA the ssgseaParam function was ignored as its execution time never passed R studio’s profiler threshold for detection (20ms), and only the call to the gsva function was considered.

## Software availability

SiPSiC is available at Bioconductor (https://bioconductor.org/packages/SiPSiC), at GitHub (https://github.com/DanielDavis12/SiPSiC).

## Competing interest statement

The authors declare no competing interests.

## Acknowledgments

We thank Michal Rabani, Michael Berger and Mor Nitzan (HUJI) for their insightful comments on the manuscript. This work was supported by the European Research Council Horizon 2020 grant 949029 (YD), and the Israel Science Foundation grant 3650/20 (YD).

## Author contributions

D.D.: Methodology, Software, Validation, Formal analysis, Investigation, Writing, Visualization. A.W.: Investigation, Validation, Visualization. Y.D.: Conceptualization, Methodology, Validation, Investigation, Resources, Writing, Visualization, Supervision, Project administration, Funding acquisition.

## Figure legends

Supplemental Fig. S1 - **(A)** Heatmap depicting z-scores of SiPSiC scores of the 20 most differential hallmark pathways in lung adenocarcinoma. Pathways are sorted by significance of differential scores, cells in each cell group are sorted by their average z-score across all shown pathways. Pathway names mentioned in the text are painted red. **(B)** Heatmap depicting z-scores of SiPSiC scores of all differential hallmark pathways (FDR < 0.01) in glioblastoma. Pathways upregulated in each group are sorted by significance of differential scores, cells in each cell group are sorted by their average z-score across all pathways upregulated in that group. Pathway names mentioned in the text are painted red. **(C)** Heatmap depicting z-scores of SiPSiC scores of all differential hallmark pathways (FDR < 0.01) in oligodendroglioma. Pathways upregulated in each group are sorted by significance of differential scores, cells in each cell group are sorted by their average z-score across all pathways upregulated in that group. Pathway names mentioned in the text are painted red.

Supplemental Fig. S2 - **(A-C)** UMAP projections based on gene expression, SiPSiC scores, Seurat Integration, Harmony and scVI, as shown in the titles on top. **(A)** Cells were clustered by Louvain algorithm according to gene expression, SiPSiC scores, Seurat Integration, Harmony and scVI, as shown in the titles on top, UMAPs show cells colored by cluster. **(B)** Cells colored by patient identity. **(C)** Cells colored by malignant meta-module assignment.

Supplemental Fig. S3 - Violin plots showing normalized pathway scores distribution of the MYC targets pathways for the monkey CD8+ T cells, as calculated by the four different methods. Significant results (FDR < 0.01) are painted red.

## Notes

### Competing Interest Statement

The authors have declared no competing interest.

### Summary of Updates

Added comparisons to VAM and ssGSEA. Added analysis of lung adenocarcinoma and oligodendroglioma. Added analysis of execution times. Added analysis of robustness to different settings of tau.

