## Supplementary figures and images for "Accurately estimating pathway activity in single cells for clustering and differential analysis"

### Supplemental Figure S1

Supplemental Fig. S1

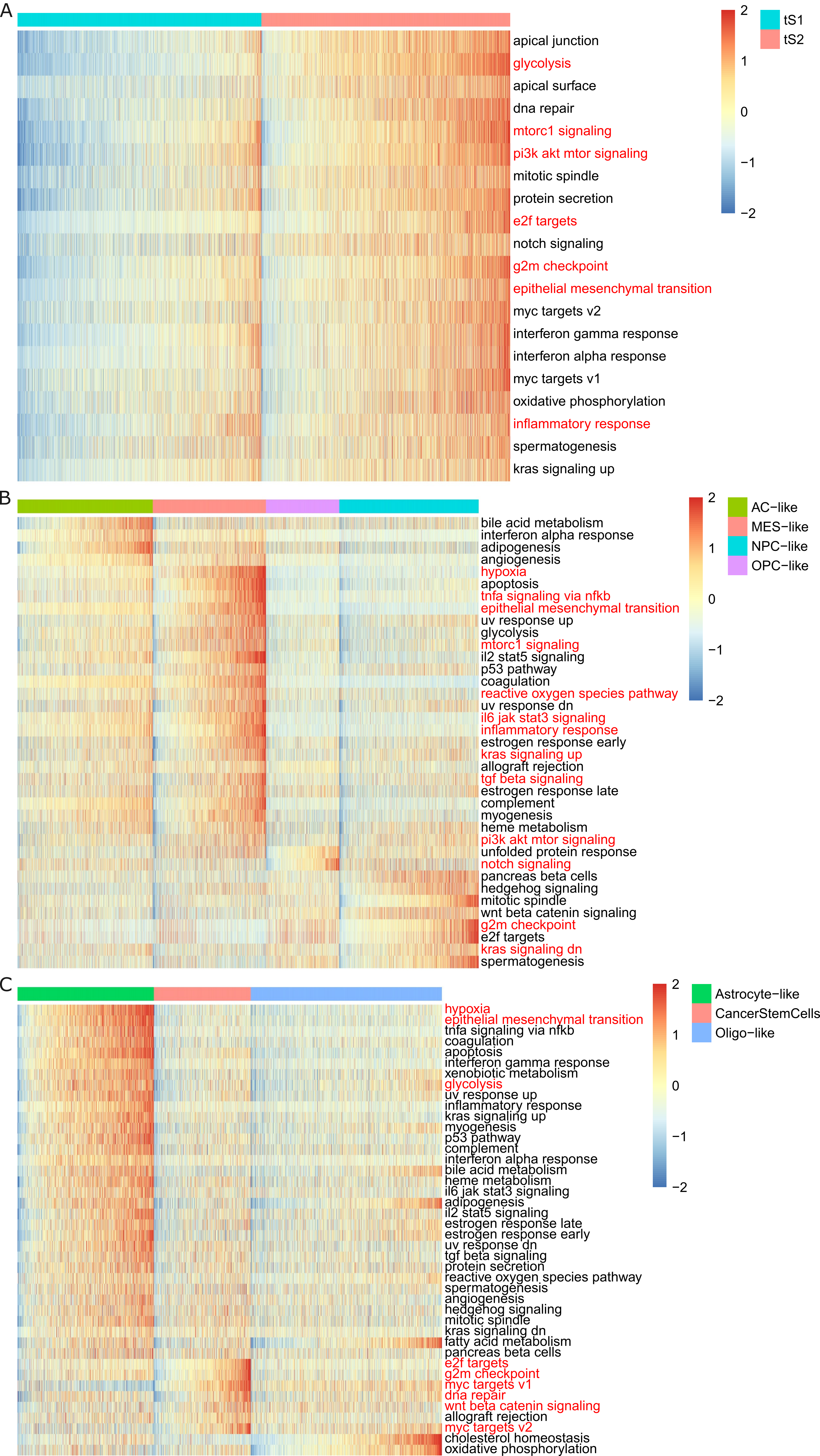

### Supplemental Figure S2

Supplemental Fig. S2

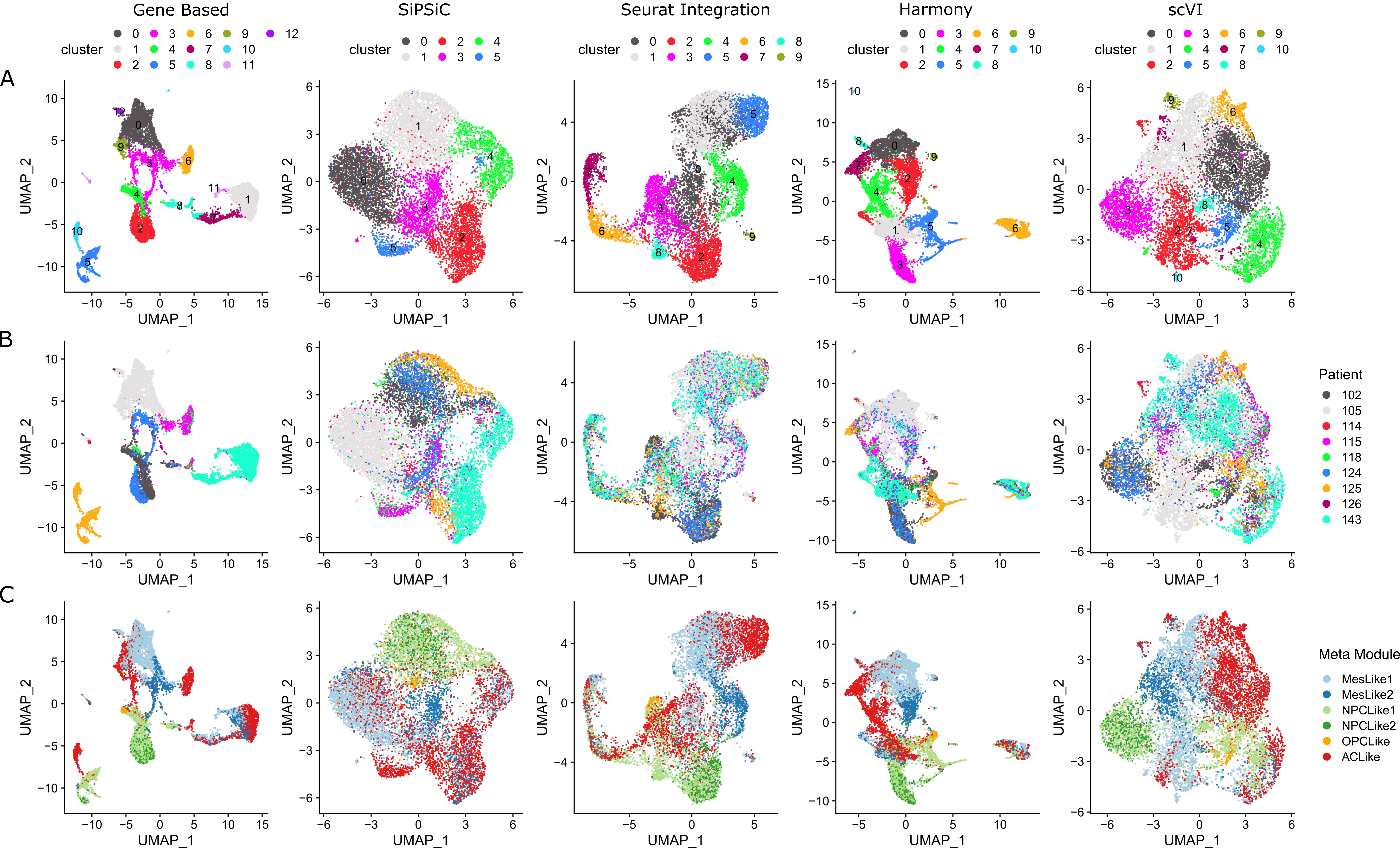

### Supplemental Figure S3

Supplemental Fig. S3

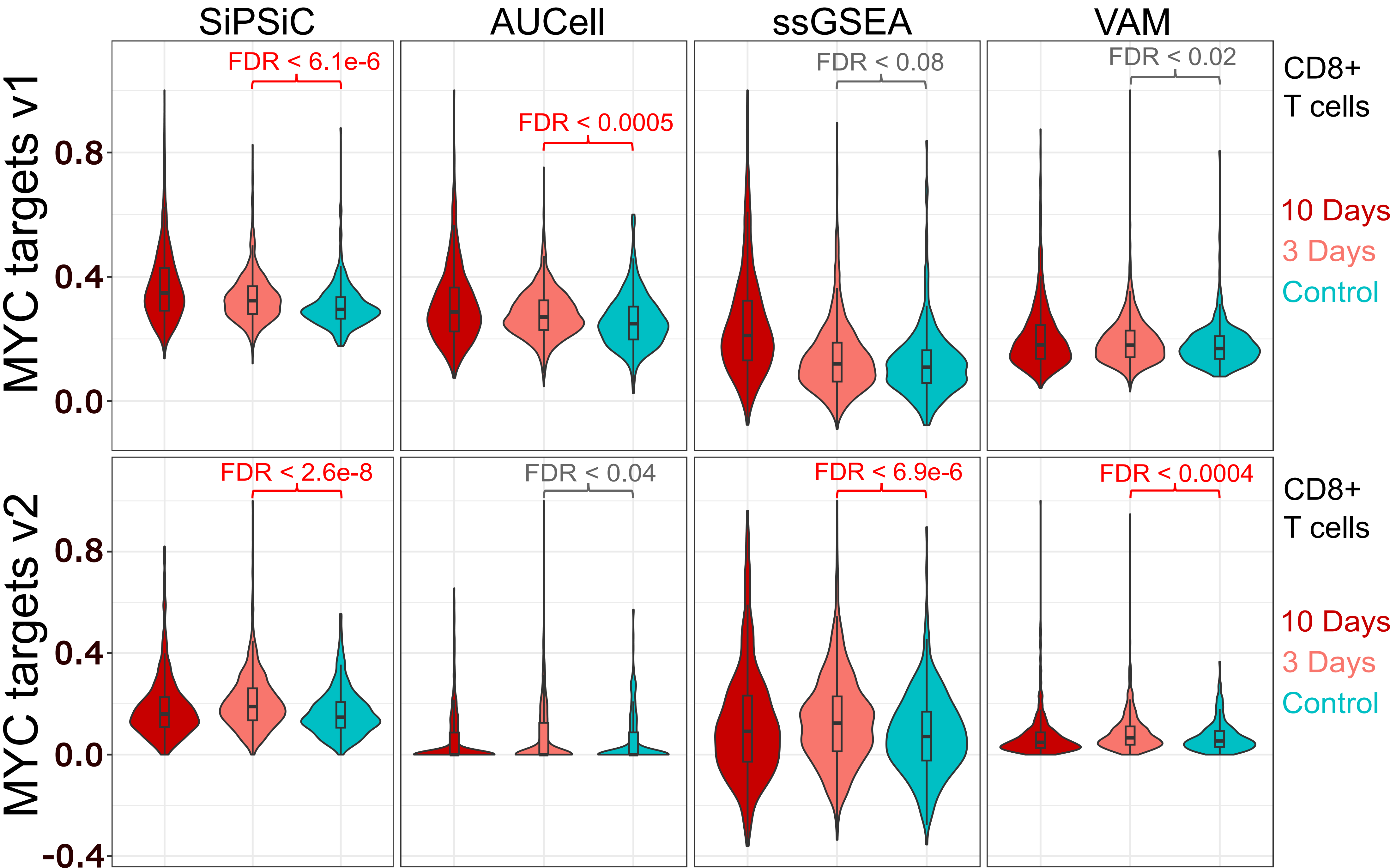
